# TEMI: Tissue Expansion Mass Spectrometry Imaging

**DOI:** 10.1101/2025.02.22.639343

**Authors:** Hua Zhang, Lang Ding, Amy Hu, Xudong Shi, Penghsuan Huang, Haiyan Lu, Paul W. Tillberg, Meng C. Wang, Lingjun Li

## Abstract

The spatial distribution of diverse biomolecules in multicellular organisms is essential for their physiological functions. High-throughput *in situ* mapping of biomolecules is crucial for both basic and medical research, and requires high scanning speed, spatial resolution, and chemical sensitivity. Here, we developed a Tissue Expansion method compatible with matrix-assisted laser desorption/ionization Mass spectrometry Imaging (TEMI). TEMI reaches single-cell spatial resolution without sacrificing voxel throughput and enables the profiling of hundreds of biomolecules, including lipids, metabolites, peptides (proteins), and N-glycans. Using TEMI, we mapped the spatial distribution of biomolecules across various mammalian tissues and uncovered metabolic heterogeneity in tumors. TEMI can be easily adapted and broadly applied in biological and medical research, to advance spatial multi-omics profiling.

## Main

Multicellular organisms consist of diverse molecular species, including nucleic acids, proteins, lipids, carbohydrates, and metabolites. These molecules actively contribute to organismal physiology and pathology, and their spatial distribution within tissues and across different cell types plays a crucial role in their functions. Advances in next-generational sequencing and spatial transcriptomic profiling methods have revolutionized the high-throughput analysis of nucleic acids with molecular specificity and spatial precision^1, 2^. However, mapping the spatial distribution of other molecular species, such as lipids, metabolites, proteins, and glycans, with high spatial resolution and throughput remains technically challenging.

Mass spectrometry imaging (MSI) has emerged as a powerful technique for spatially mapping molecules within tissue samples in a label-free manner^3–7^. However, its broad application in basic and medical research has been limited by its low spatial resolution. Recent advances in instrumentation have improved MSI spatial resolution^8–12^, but their broad adoption has been limited by prohibitive costs and specialized hardware requirements. To circumvent this challenge, we have developed an easily adaptable and inexpensive tissue expansion method compatible with MSI, named TEMI.

## Results

### Development of tissue expansion method for MSI

Tissue expansion is usually conducted by chemically anchoring proteins into an *in situ*-synthesized hydrogel, followed by harsh denaturation through combinations of high temperature, detergent, and proteolysis to enable uniform swelling of the tissue-hydrogel material^13^ up to ∼ten-fold linearly upon washing with deionized water^14, 15^, or ∼20-fold with sequential embedding rounds^16^. We hypothesized that with no proteolysis, detergent, or high temperature treatment in tissue expansion, biomolecules including lipids, protein-associated metabolites, peptides, and N-glycans, would be better retained through interactions with anchored, native-state proteins, which may offer an opportunity for high-spatial resolution mapping of these biomolecules within the tissue using MSI (**Figure 1a**).

**Figure 1.**
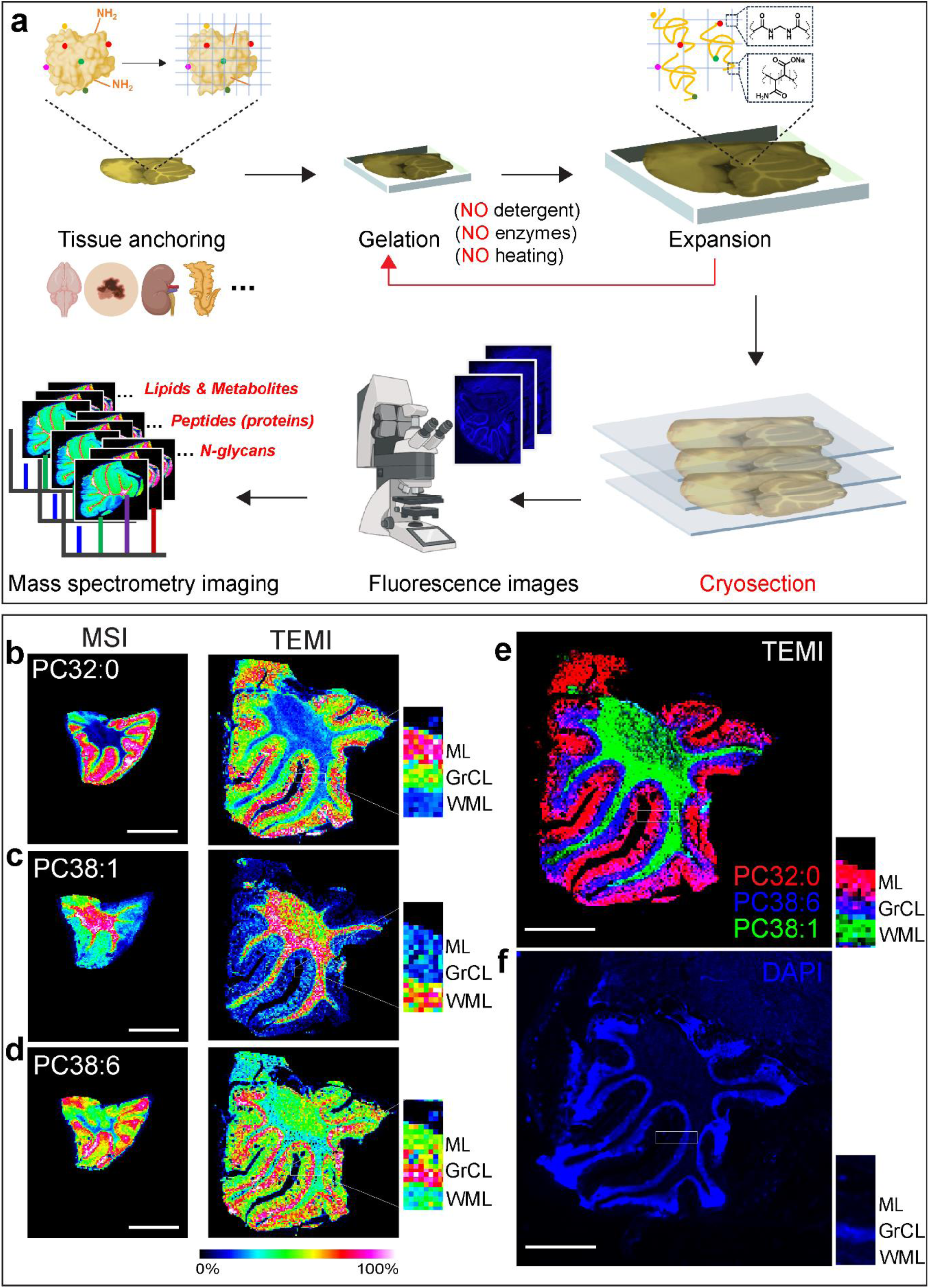
TEMI mapping of lipid distribution across cerebellar layers. a. Concept and workflow of Tissue Expansion coupled with MALDI-MSI (TEMI) for mapping biomolecules with improved spatial resolution. The color dots denote small molecule metabolites, the blue network denotes the hydrogel polymers, with chemically anchored proteins; b-e. Representative MS images of unexpanded control (left panel, MSI) and double-embedded (right panel, TEMI) cerebellum using a 50 *µ*m laser beam raster scanning, with lipid species specifically enriched in three distinctive layers: b. PC (32:0) ([M + H]^+^, *m/z* 734.568), molecular layer (ML); c. PC (38:1) ([M + H]^+^, *m/z* 816.653), white matter layer (WML); d. PC (38:6) ([M + H]^+^, *m/z* 806.573), granular cell layer (GrCL); e. Overlay of TEMI images of PC (32:0) (red), PC (38:6) (green), and PC (38:1) (blue), highlighting the functional layers of cerebellum; f. DAPI staining fluorescent image of the expanded cerebellum tissue. All the MSI images obtained with mass error tolerance of 10 ppm. The scale bar is 2 mm.

We started by optimizing cryosectioning conditions for expansion-embedded tissue, as we hypothesized that an uneven layer of hydrogel on the tissue surface could interfere with MALDI laser penetrance and ion release. We used mouse cerebellar tissue embedded with a high monomer, high toughness gel^15^. We explored different sucrose concentrations and slice thicknesses, and found that 30% sucrose provided the best performance when cryosectioning the expanded tissue-gel material. At other sucrose concentration, we did not obtain useful slices for conducting MSI. Obtaining intact tissue slices thinner than 30 *µ*m became increasingly difficult (**Supplementary Figure 1**). MALDI-MS spotting of these tissue sections indicated that the signal intensities exhibit no significant decline as the thickness of tissue sections increases from 10 to 30 *µ*m (**Supplementary Figure 2**).

Conventional tissue-expansion methods that use high temperature or proteolysis have recently been coupled with MSI^17–19^. However, we were concerned that these harsh treatments could compromise performance by causing both signal loss and signal delocalization^17^. To compare overall analyte retention, we extracted lipids from tissue sections expanded with and without proteinase K treatment, as well as unexpanded control sections, and then analyzed them using HPLC-ESI-MS/MS. Overall, the detected lipid species were only modestly reduced in the tissue expanded without proteinase K digestion compared to the control, while the tissue expanded with digestion exhibited strong signal loss (**Supplementary Figure 3 and Supplementary Figure 4**). We also expanded mouse brain cerebella treated with proteinase K or trypsin digestion according to published methods^18, 19^, and only a few slices survived after drying before MSI (**Supplementary Figure 5, Supplementary Note 1**). When running MSI, we noticed that without post-gel cryosectioning (as described in recently reported methods^18, 19^), in practice, the expanded tissue was sometimes located beneath an uneven layer of hydrogel, which prevented the MALDI laser beam from efficiently releasing material from the tissue. Moreover, the presence of hydrogel on the tissue surface interfered with the crystallization of analytes and the MALDI matrix, significantly impacting MSI results.

When expanding the tissue without denaturation, we found that full expansion in deionized water typically results in cracks or macroscale gel deformation. Thus, we started by limiting the expansion extent to ∼1.5-fold by swelling in 1× PBS. Strikingly, even with ∼1.5-fold expansion, MSI via a 50 *µ*m laser raster scan showed more structural features compared with unexpanded controls (**Supplementary Figure 6**). These results confirm the effectiveness of denaturation-free expansion in revealing chemical distribution with enhanced spatial resolution. They also suggest that TEMI makes it possible to conduct tissue scanning with both high spatial resolution and high voxel throughput. In unexpanded samples, achieving the same effective spatial resolution requires a finer laser beam for raster scanning, which in turn increases acquisition time.

We next reasoned that re-embedding the gelled tissue with another round of gelation might produce an interpenetrating gel network with the toughness required to expand non-denatured tissue isotropically down to the micron-scale features required for this application. After two rounds of gel embedding, mouse cerebellar slices expanded ∼2.5-fold in 1× PBS. We observed that the expanded gels had some residual curvature but could easily be laid flat prior to freezing without macroscopic tissue distortion (**Extended Data Figure 1a-d, Supplementary Note 2**). Expanded cerebellar tissue sections with anatomically matched unexpanded controls were imaged with 1,5-diaminonaphthalene (DAN) matrix. MALDI-MS spotting experiments revealed similar mass spectral patterns of lipid molecules from the corresponding areas on the control and expanded tissue slides in both positive and negative ionization modes (**Extended Data Figure 2**). The expanded samples showed a clean background without contaminant signals from the hydrogel material (**Supplementary Figure 7**). It was observed that MS signal intensities were decreased slightly in the expanded tissue samples, which may be attributed to smaller effective voxels necessarily containing fewer analytes. In addition, targeted lipid signals were exclusively detected within the expanded tissue area, with no corresponding signal observed in the adjacent blank hydrogel area (**Supplementary Figure 8**), indicating a lack of chemical delocalization in the expansion sample.

### Lipid distribution specificity across cerebellar layers via TEMI

Non-denatured, ∼2.5-fold expanded cerebellar sections were next imaged with a 50 *µ*m raster laser scan with dual-polarity ionization on the same tissue, for an effective lateral resolution of ∼20 *µ*m. Strikingly, the molecular layer (ML), granular cell layer (GrCL), and white matter layer (WML) were more clearly distinguished in each cerebellar folium compared to the unexpanded control sample (**Figure 1b-f, Supplementary Figures 9 and 10**). Furthermore, leveraging this enhanced spatial resolution, we were able to identify lipid molecules that were specifically enriched in different functional layers, such as PC (32:0), PC (38:1), and PC (38:6) for ML, GrCL, and WML, respectively (**Figure 1b-d**). This enables us to reveal the spatial biomolecular heterogeneity of cerebellum in conjunction with its structural organization (**Figure 1e and f**) and uncover previously unknown lipid fingerprints for each functional layer (**Supplementary Figures 9-12, Supplementary Tables 1 and 2**).

We further explored the potential of TEMI in improving spatial resolution. Cerebellar slices were expanded with three rounds of embedding without denaturation, for a final expansion factor of ∼3.5-fold. Imaging these ∼3.5-fold expanded tissue sections with a 10 *µ*m raster laser scan, or a ∼2.9 *µ*m effective scanning resolution, revealed additional details compared with ∼2.5-fold expansion (**Figure 2 and Extended Data Figure 3**). At this resolution, individual Purkinje cells became visible (**Figure 2**), revealing an enrichment of specific lipid species, such as PC (38:1), PC (40:5), PC (40:6), and the others, while the lipid species, such as PC (32:0) and PC (34:1) were absent in these cells (**Supplementary Table 3**). In contrast, the individual Purkinje cells were barely distinguishable in the unexpanded control sample when using conventional MSI with the same laser beam scan (**Extended Data Figure 3**).

**Figure 2.**
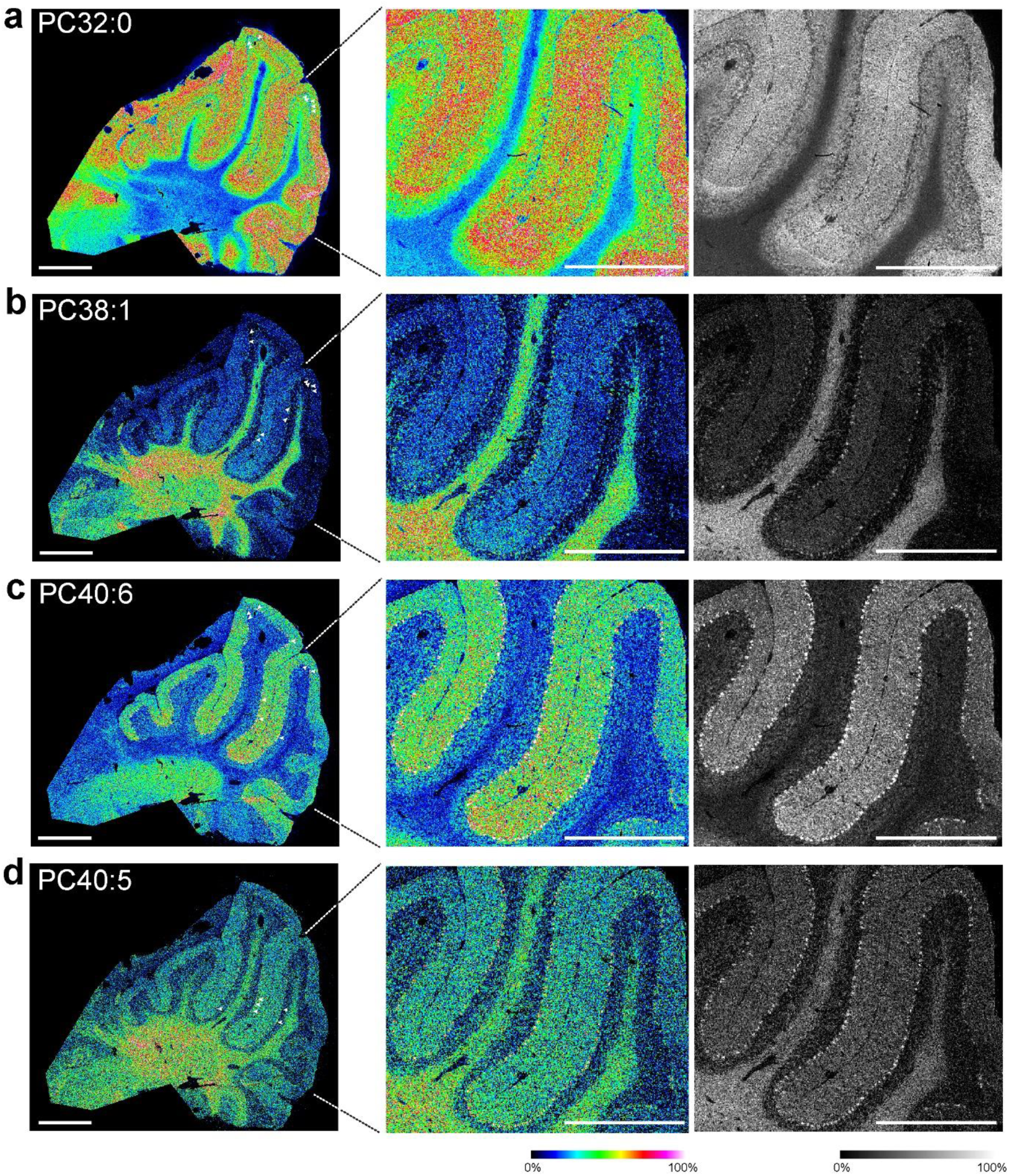
Single-cell resolution TEMI mapping of Purkinje cells. a-d. Representative TEMI images of a ∼3.5-fold linearly expanded mouse cerebellum tissue through a 10 *µ*m laser beam raster scanning under positive mode: a. PC (32:0) ([M + H]^+^, *m/z* 734.5747), b. PC (38:1) ([M + H]^+^, *m/z* 816.647), c. PC (40:6) ([M + H]^+^, *m/z* 834.603), d. PC (40:5) ([M + H]^+^, *m/z* 836.617). All the MSI images obtained with mass error tolerance of 10 ppm. Arrow heads highlight several Purkinje cells as examples. The scale bar is 2 mm.

### TEMI profiling of metabolite distribution in cerebella

In addition to glycolipids and phospholipids, we also imaged metabolites in the mouse cerebella expanded ∼2.5-fold linearly. We applied 2,5-Dihydroxybenzoic acid (DHB) and N-(1-Naphthyl) ethylenediamine dihydrochloride (NEDC) matrix for positive and negative modes respectively (**Figure 3a**). Among 187 metabolite features detected, some exhibited distinctive spatial organization among ML, GrCL, and WML, which were not discernible in the unexpanded controls (**Figure 3b-g and Supplementary Figures 13 and 14**). This result further supports the spatial metabolic heterogeneity of cerebellum in conjunction with its structural organization. While we were able to annotate some metabolites with their chemical identities, such as choline, hexose-phosphate, oleoylethanolamide (OEA), and 2-arachidonoylglycerol (2-AG) (**Supplementary Figure 15 and Extended Data Figure 4**), fully and accurately annotating all detected metabolites is difficult due to the lack of a reference library based on MALDI-MSI.

**Figure 3.**
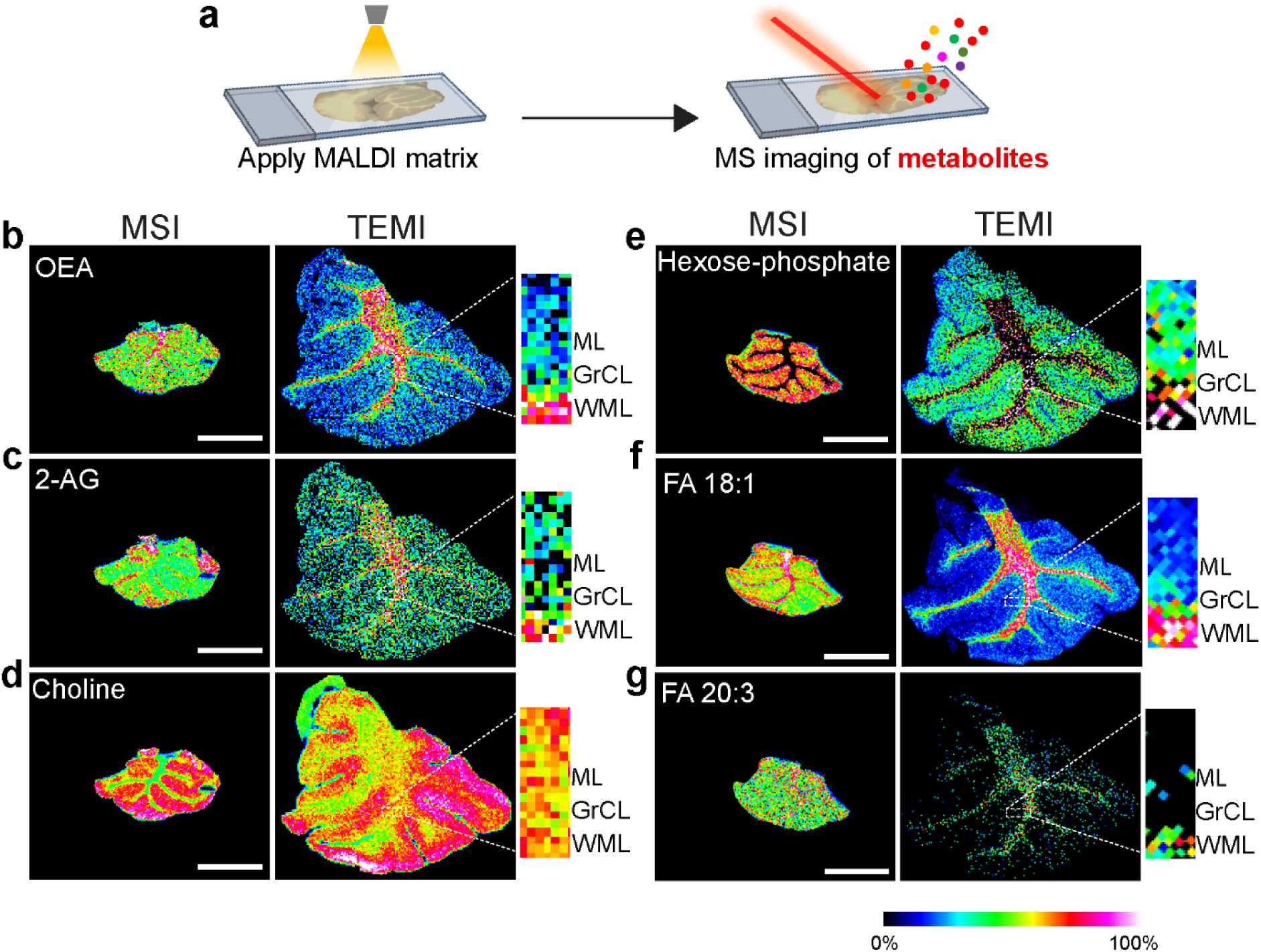
TEMI mapping of small metabolites in mouse cerebellum. a. Schematic workflow for detecting metabolites in imaging mass spectrometry. b-g. Representative MS images of the unexpanded control (left panel, MSI) and double-embedded (right panel, TEMI) cerebellum under positive (b-d) and negative (e-g) ionization mode with these small molecules specifically enriched in the three distinctive layers of ML, GrCL, and WML. b. Oleoyl ethanolamide (OEA, [M -H_2_O+ H]^+^, *m/z* 308.296), WML; c. 2-Arachidonoyl glycerol (2-AG, [M-H_2_O + H]^+^, *m/z* 361.277), WML; d. Choline ([M + H]^+^, *m/z* 104.107), ML; e. Hexose-phosphate ([M - H]^−^, *m/z* 259.022), ML; f. Fatty acid 18:1 (FA18:1, [M - H]^−^, *m/z* 284.249), WML; g. Fatty acid 20:3 (FA 20:3, [M - H]^−^, *m/z* 305.248), WML. The MSI experiments were carried out using a 50 *µ*m laser raster scanning and the MSI images obtained with mass error tolerance of 10 ppm. The scale bar is 2 mm.

### Mapping cerebellar proteins and N-glycans with TEMI

We also imaged peptides, proteins, and N-glycans using TEMI in the mouse cerebella expanded ∼3.5-fold linearly. For imaging proteins, we have applied *in situ* tryptic digestion of both unexpanded and expanded tissue sections to release peptides for MSI and LC-MS proteomic profiling. Based on the LC-MS proteomic results and *in situ* MALDI-MS/MS (**Supplementary Table S4 and Extended Data Figure 5**), we matched 57 features within 5 ppm mass tolerance, including peptides from myelin basic protein group (MBP), histone H2B protein group (H2B), as well as other proteins (**Figure 4b-d, Supplementary Figure 16**). Interestingly, these proteins exhibited distinctive distributions, with histone H2B protein groups enriched in GrCL layer, while MBP enriched in WML.

**Figure 4.**
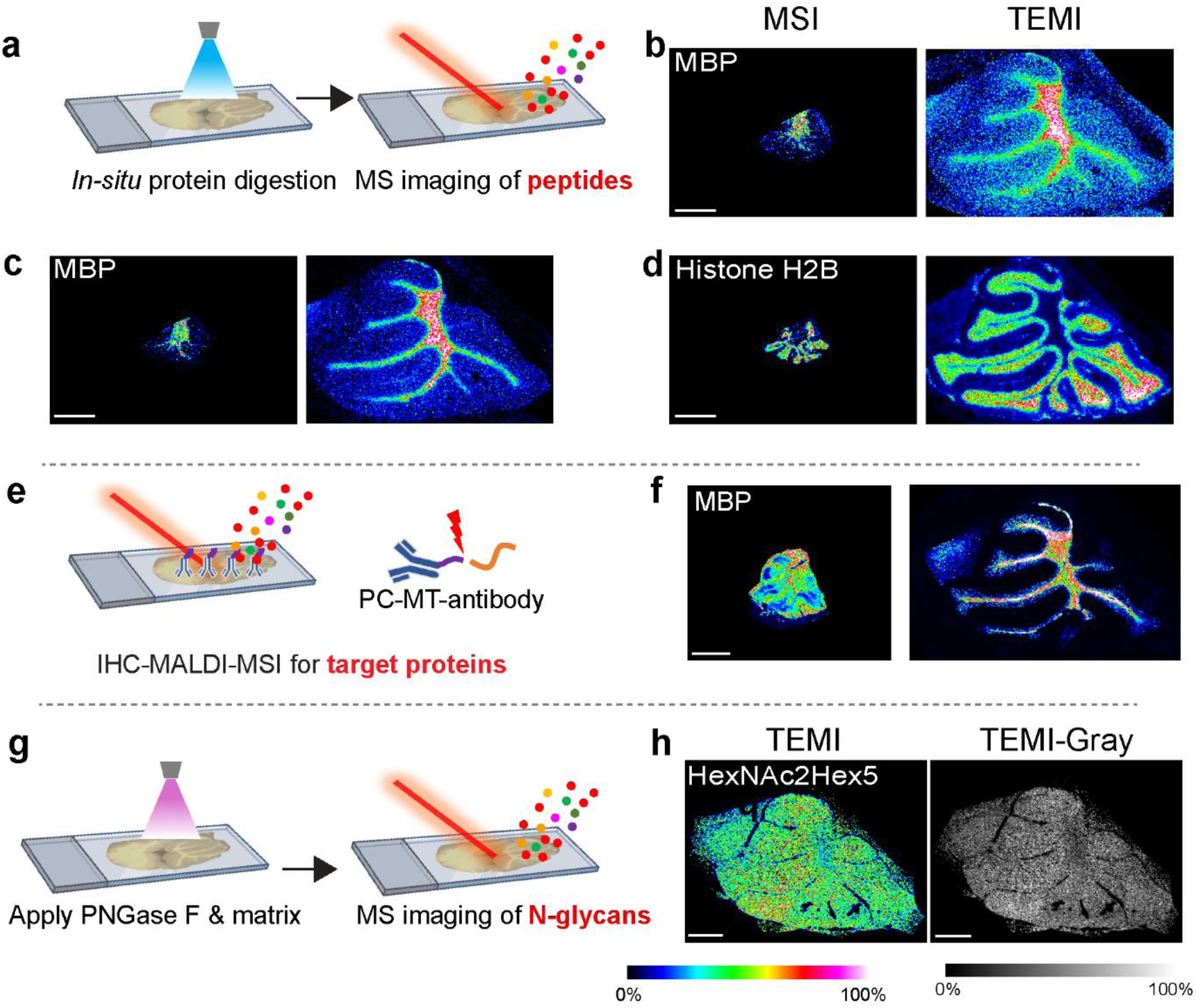
TEMI mapping of proteins and N-glycans in mouse cerebellum. a. Schematic of MSI tryptic peptides using *in-situ* trypsin digestion of expanded tissue sections; b. Peptide sequence: TTHYGSLPQKSQHGR from myelin basic protein group ([M + H]^+^, *m/z* 1696.856); c. Peptide sequence: HRDTGILDSIGRFFSGDRGAPK from myelin basic protein group ([M + H]^+^, *m/z* 2402.242); d. Peptide sequence: E[+14]VQTAVRLLLPGELAK from histone H2B protein group ([M + H]^+^, *m/z* 1751.045); e. Schematic workflow for detecting proteins using IHC-MALDI-MSI. f. Distribution of myelin basic protein in the unexpanded and TEMI samples revealed by IHC-MALDI-MSI ([M + H]^+^, *m/z* 1366.042); g. Schematic of *in situ* N-glycans release using PNGase F for MSI imaging; h. TEMI ion image of N-glycan HexNAc2Hex5 ([M + H]^+^, *m/z* 1257.429). The MSI images obtained with mass error tolerance of 10 ppm. The scale bar is 2 mm.

Furthermore, we utilized immunohistochemistry (IHC) photocleavable mass-tag (PC-MT) antibodies on both unexpanded and TEMI samples (**Figure 4e**). IHC-MALDI-MSI confirmed the distinctive pattern of MBP, as well as other proteins revealed by bottom-up tryptic digestion in the TEMI sample (**Figure 4f, Supplementary Figure 17**). In parallel, we successfully detected up to 29 N-glycans in both unexpanded and TEMI samples (**Figure 4g–h, Supplementary Figure 18, and Supplementary Table 5**). Although most N-glycans are ubiquitously distributed, TEMI revealed that certain N-glycan, such as HexNAc2Hex5, exhibited a moderate degree of regional distinctiveness (**Figure 4h**).

### Revealing high metabolic heterogeneity in tumors with TEMI

In addition to cerebellum, we also tested the applicability of TEMI to *in situ* biomolecular mapping of cancer tissues. Tumor tissue from a murine melanoma model was subjected to a 3-round gelation-expansion cycle, resulting in a ∼3.5-fold linear expansion. With a consistent 50 *µ*m of laser raster scanning, the expanded tumor tissue revealed more detailed features compared to the control unexpanded sample (**Figure 5a-d**). The enhanced resolution enabled the spatial separation of metabolic features, such as phospholipids PC (36:1) and PC (38:4), which both had less distinct spatial patterns without expansion (**Figure 5c-d**). The tumor tissue exhibits high heterogeneity as revealed from spatial distribution of the lipids. Based on spectra clustering using Bisecting k-means, we were able to segment the tumor tissue into 21 dominant regions using the TEMI method, while the unexpanded tissue was only able to segment into 3 dominant regions when we set the minimum cluster of 600 spectra (**Figure 5e-f**). These results support the ability of TEMI to uncover metabolic heterogeneity in tumor tissues.

**Figure 5.**
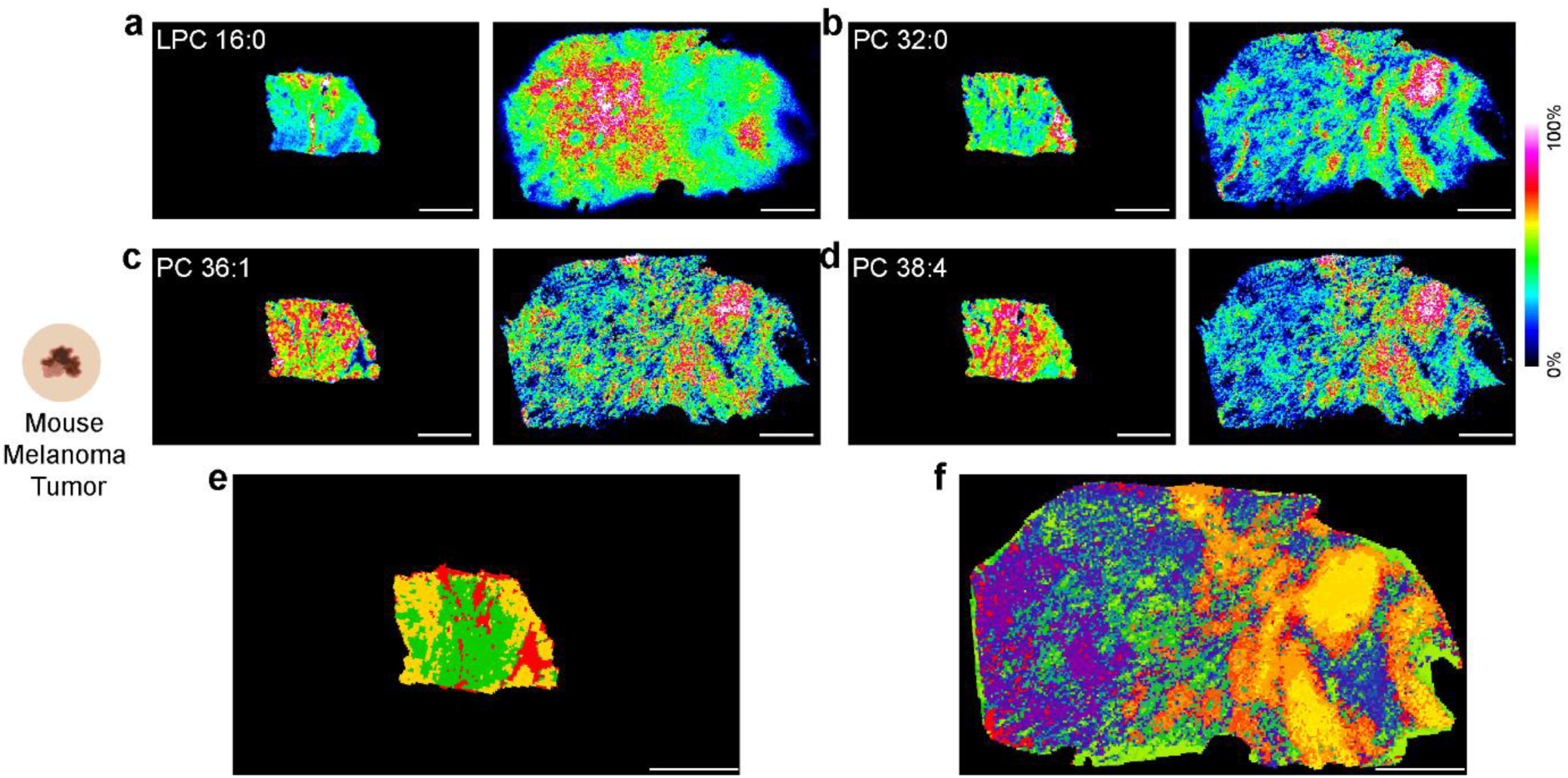
Revealing metabolic heterogeneity in mouse tumor with TEMI. a-f. MS images of representative lipid species detected from unexpanded control (MSI, left panel) and expanded tumor tissue (TEMI, right panel), including a. LPC (16:0) ([M + H]^+^, *m/z* 496.340), b. PC (32:0) ([M + H]^+^, *m/z* 734.568), c. PC (36:1) ([M + H]^+^, *m/z* 788.614), d. PC (38:4) ([M + H]^+^, *m/z* 810.598), e and f, Spectra clustering segmentation of the tumor tissue into 3 dominant regions in MSI control and 21 dominant regions in the TEMI sample. The MSI experiments were carried out using a 50 *µ*m laser beam raster scanning and the MSI images obtained with mass error tolerance of 10 ppm. The scale bar is 2 mm.

### Broad application of TEMI in organ imaging

Furthermore, we have applied TEMI for biomolecular mapping of other organs including mouse kidney and pancreas (**Figure 6**) and revealed the spatial distribution of different lipid species within those different tissues. Together, these results support broad application of TEMI for spatial molecular mapping. Furthermore, at each stage of gel embedding, we measured the tissue deformations in the cerebellum, kidney, and pancreas, finding that the maximum measurement errors were less than 12% (**Extended Data Figure 1, Supplementary Table 6**).

**Figure 6.**
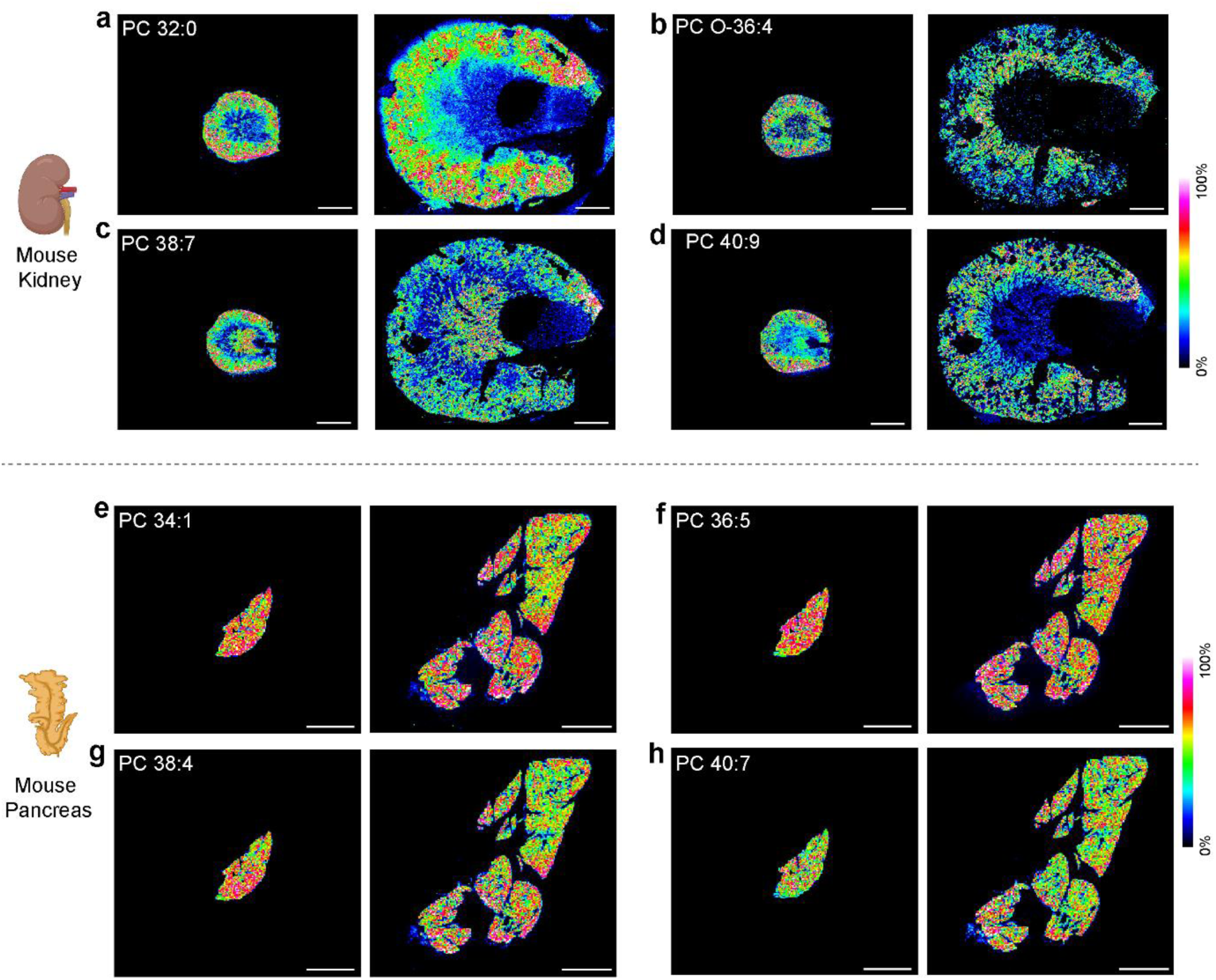
TEMI-based metabolic mapping of kidney and pancreas. a-d. MS images of representative lipid species detected from unexpanded control (MSI, left panel) and expanded mouse kidney tissue (TEMI, right panel), including a. PC (32:0) ([M + H]^+^, *m/z* 734.568), b. PC (O-36:4) ([M + H]^+^, *m/z* 768.586), c. PC (38:7) ([M + H]^+^, *m/z* 804.549), d. PC (40:9) ([M + H]^+^, *m/z* 828.550). e-h. MS images of representative lipid species detected from unexpanded control (MSI, left panel) and expanded mouse pancreas tissue (TEMI, right panel), including e. PC (34:1) ([M + H]^+^, *m/z* 760.583), f. PC (36:5) ([M + H]^+^, *m/z* 780.551), g. PC (38:4) ([M + H]^+^, *m/z* 810.600), h. PC (40:7) ([M + H]^+^, *m/z* 832.581). The MSI experiments were carried out using a 50 *µ*m laser beam raster scanning and the MSI images obtained with mass error tolerance of 10 ppm. The scale bar is 2 mm.

## Discussion

Together, this work demonstrates the TEMI method for spatially resolved omics in intact tissues at high spatial resolution and chemical sensitivity, thereby revealing *in situ* biomolecular heterogeneity. In this proof-of-concept study, TEMI was applied not only to lipidomics and metabolomics but also to the systematic profiling of proteins and N-glycans. Recently, three published and two preprint studies have utilized tissue expansion protocols with varying harsh denaturing treatments including high temperature, detergent, proteinase K, or trypsin digestion followed by MALDI-MSI^17–21^. These published methods overlooked the importance of preserving non-denatured tissue during tissue expansion or the essential step of cryosectioning following the expansion, which we have found to be important for robust analyte detection. Harsh denaturing treatments can reduce signals, making detection sensitivity a limiting factor for method performance, or even lead to severe biomolecule delocalization upon expansion^17^. In contrast, the new multi-embedding expansion protocol used in the TEMI approach presented here achieves expansion without denaturation, enabling the preservation of biomolecules in expanded tissues and revealing their distribution with demonstrated single-cell resolution. Whereas a previously reported tissue stretching strategy using glass beads embedded in Parafilm improves MSI spatial resolution by fragmentation of the tissue at the cellular level^22^, TEMI fragments tissue at the molecular scale, is compatible with standard MSI processing and is compatible with correlative light microscopy. Therefore, for the future application of tissue expansion in MALDI-MSI, TEMI offers a promising and crucial platform to uncover the spatial and chemical specificity of various biomolecules and reveal biomolecular heterogeneity across diverse biomedical samples.

## Acknowledgments

Aspects of this work were supported in part by National Institutes of Health (NIH) grants R01 AG078794 (L.L.), R01 DK071801 (L.L.), R01 AG052324 (L.L.), P01 CA250972 (L.L.), DP1DK113644 (M.C.W.) and the United States Department of Agriculture (2018-67001-28266) (L.L.). M.C.W. and P.W.T. were supported by the Howard Hughes Medical Institute (HHMI). The timsTOF fleX MALDI-2 mass spectrometer was acquired using NIH shared instrument grant S10 OD028473 (L.L.). H.Z. and H.L. thank the funding support for a Postdoctoral Career Development Award provided by the American Society for Mass Spectrometry. L.L. would like to acknowledge a Pancreas Cancer Pilot grant from the University of Wisconsin Carbone Cancer Center (233-AAI9632), a Diabetes Research Center (DRC) pilot and feasibility grant from Washington University/University of Wisconsin-Madison (P30 DK020579), NIH shared instrument grants S10 RR029531 (L.L.) and S10 OD025084 (L.L.), Research Forward grant support provided by the University of Wisconsin-Madison Office of the Vice Chancellor for Research with funding from the Wisconsin Alumni Research Foundation, as well as a Vilas Distinguished Achievement Professorship and Charles Melbourne Johnson Distinguished Chair Professorship, with funding provided by the Wisconsin Alumni Research Foundation and the University of Wisconsin-Madison School of Pharmacy. We thank Dr. Zachary Morris from the Department of Human Oncology, UW-Madison, for providing the mouse melanoma tumor samples. We thank Dr. Gargey B. Yagnik and Dr. Mark J. Lim from AmberGen for their support with the PC-MT-IHC-MALDI-MSI experiments. Tissue schematic diagram created in BioRender: Ding, L. (2025) https://BioRender.com/l96l933. We appreciate Dr. Yue Zhou for providing protocol, Dr. Nan Wang for maintaining the Orbitrap Fusion Lumos and Orbitrap Ascend Mass Spectrometers (Janelia Research Campus, HHMI), and Dr. Jiefu Li, Malik Ebbini, Kelly H Lu for critical reading of the manuscript.

## Author Information

These authors contributed equally: Hua Zhang & Lang Ding.

These authors jointly supervised this work: Paul W. Tillberg, Meng C. Wang & Lingjun Li

## Author Contributions

H.Z., L.D., P.W.T, M.C.W., and L.L. conceptualized the study and experimental design. H.Z., L.D., A.H., P.H., H.L., and X.S. conducted experiments. H.Z., L.D., P.W.T, M.C.W., and L.L. analyzed the data. H.Z., L.D., P.W.T, M.C.W., and L.L. wrote the paper, and all authors provided editorial feedback.

## Associated Content

**Supplementary Information** is available for this paper.

Additional information as noted in text.

---

**Extended Data Figure 1.**
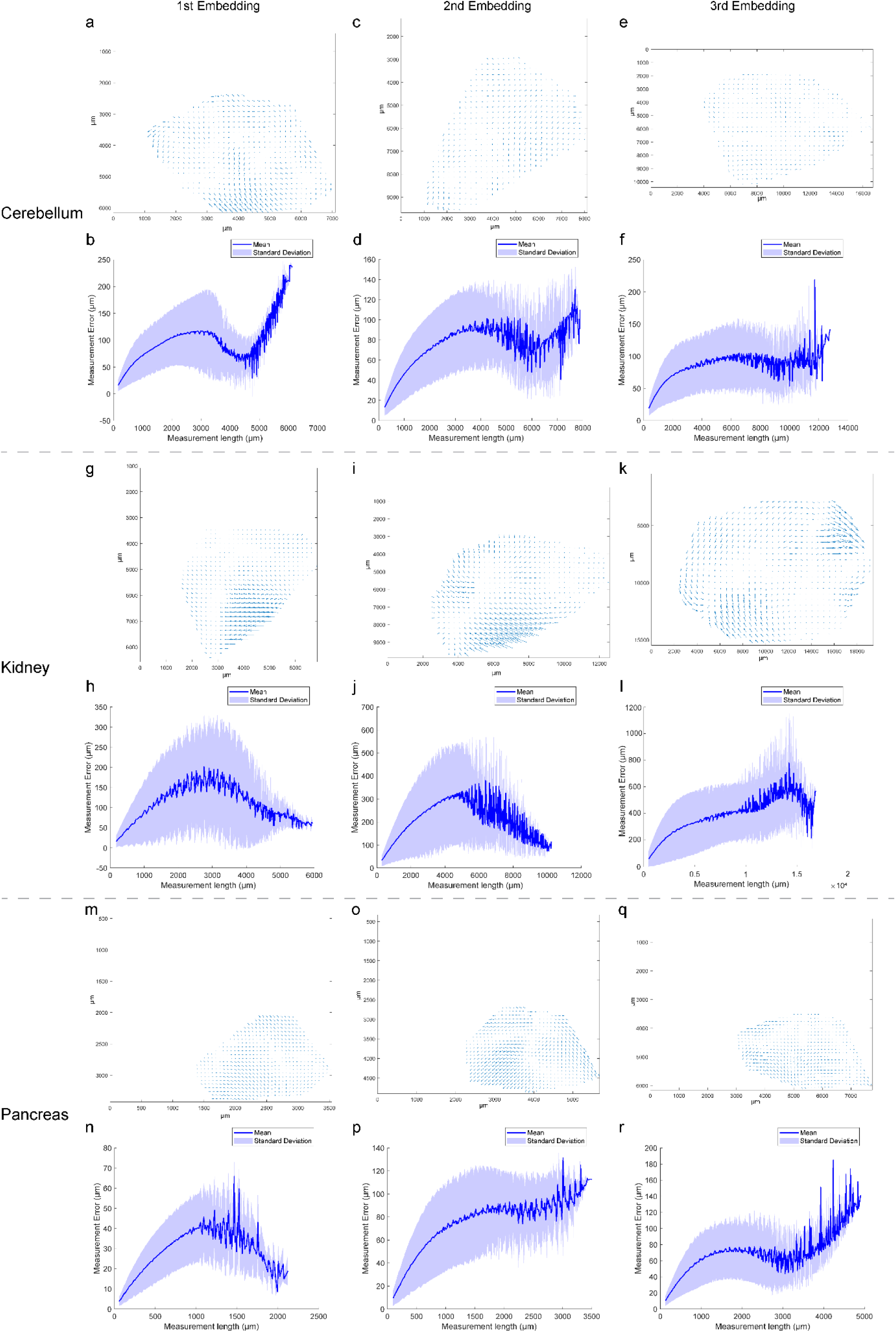
Deformation maps and quantification of measurement errors between the 1st, or 2nd, or 3rd embedding and pre-expansion of different mouse tissue types. Deformation map between: a. the 1st embedding and pre-expansion. c. the 2nd embedding and pre-expansion. e. the 3rd embedding and pre-expansion of cerebellum. g. the 1st embedding and pre-expansion. i. the 2nd embedding and pre-expansion. k. the 3rd embedding and pre-expansion of kidney. m. the 1st embedding and pre-expansion. o. the 2nd embedding and pre-expansion. q. the 3rd embedding and pre-expansion of pancreas. Quantification of measurement error (y-axis, deviation from expected distance assuming isotropic expansion) between pairs of points between: b. the 1st embedding and pre-expansion. d. the 2nd embedding and pre-expansion. f. the 3rd embedding and pre-expansion of cerebellum. h. the 1st embedding and pre-expansion, j. the 2nd embedding and pre-expansion, l. the 3rd embedding and pre-expansion of kidney. n. the 1st embedding and pre-expansion. p. the 2nd embedding and pre-expansion. r. the 3rd embedding and pre-expansion of pancreas for a measurement length in the expanded coordinates (x-axis).

**Extended Data Figure 2.**
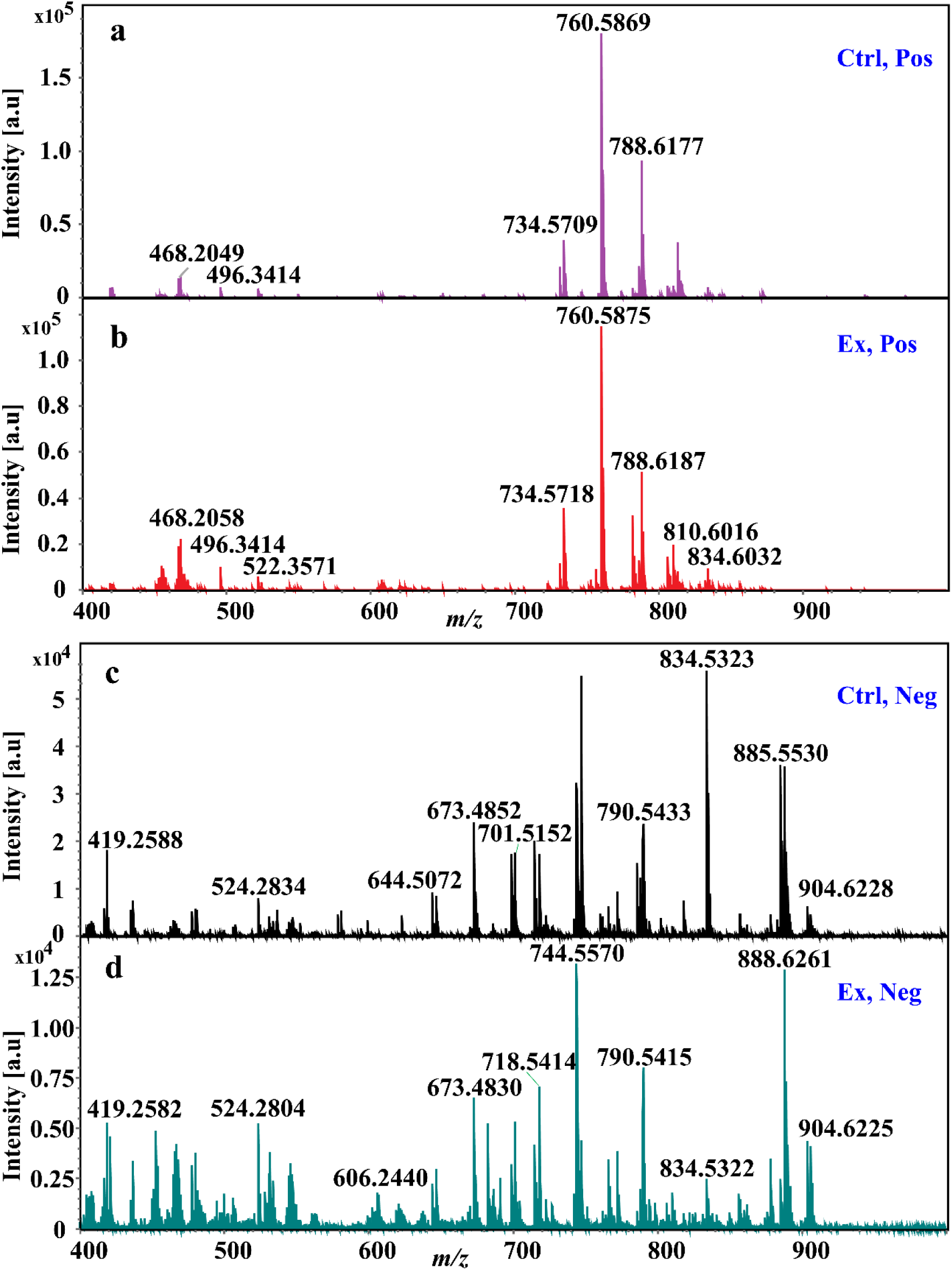
MALDI-MS spotting mass spectra from the corresponding area on control (Ctrl, unexpanded) and expanded (Ex, ∼2.5-fold linear) mouse brain cerebellar tissue under positive (Pos) and negative (Neg) ionization mode: a. mass spectra of matched control tissue under positive mode; b. mass spectra of expanded tissue under positive mode; c. mass spectra of adjacent control tissue under negative mode; b. mass spectra of expanded tissue under negative mode.

**Extended Data Figure 3.**
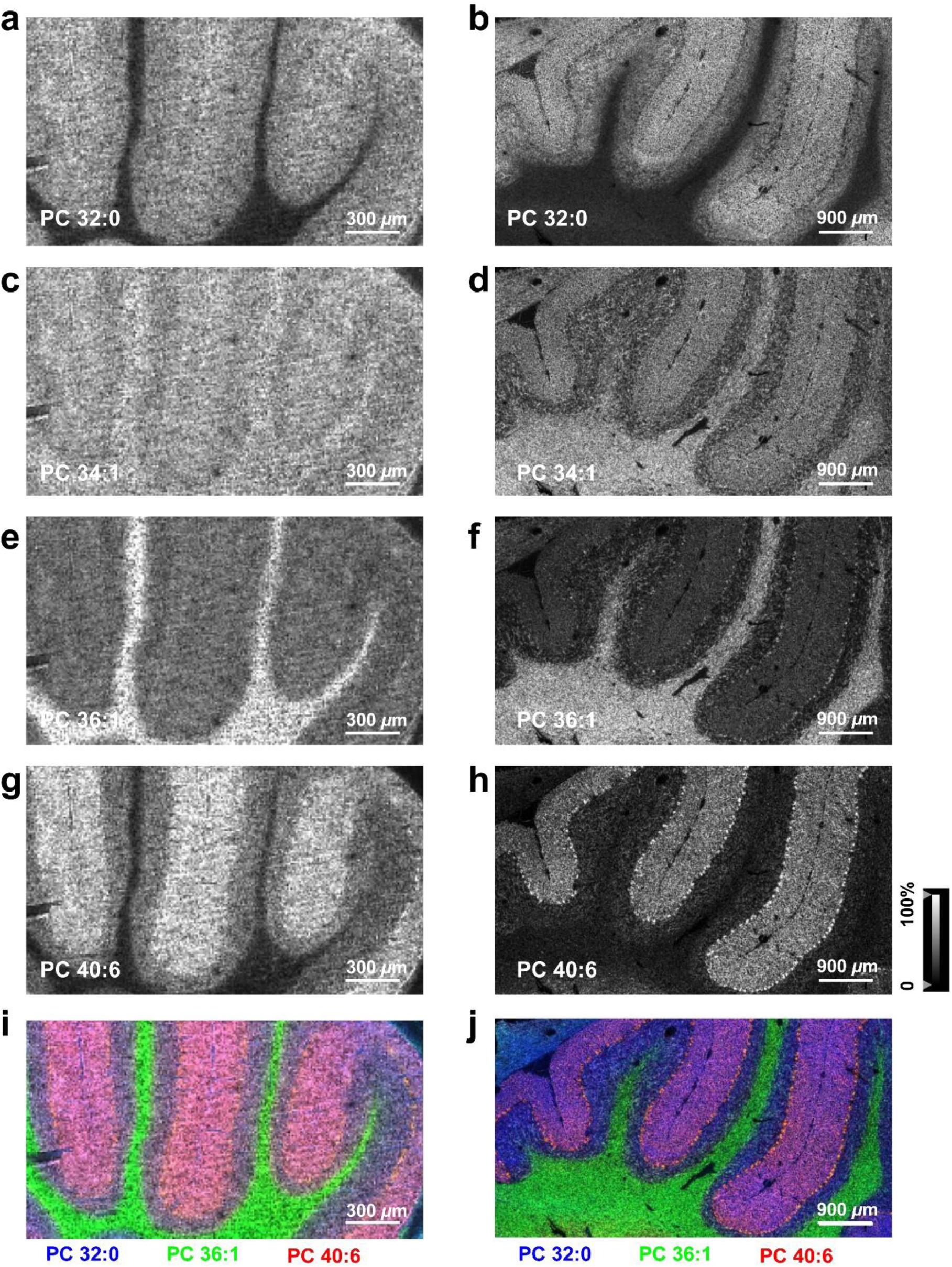
High spatial resolution TEMI of mouse cerebellum after 3 cycles of gel-embedding-expansion. The experiments were carried out using a 10 *µ*m laser beam raster scanning for both unexpanded control tissue (left: a, c, e, g, and i) and expansion treated tissue (right: b, d, f, h, and j), including a and b. For PC (32:0) ([M + H]^+^, *m/z* 734.575); c and d. For PC (34:1) ([M + H]^+^, *m/z* 760.587); e and f. For PC (36:1) ([M + H]^+^, *m/z* 788.621); g and h. For PC (40:6) ([M + H]^+^, *m/z* 834.603); overlapped ion images of PC (32:0) (in blue), PC (36:1) (in green), and PC (40:6) (in red) from the unexpanded control (i) and 3 cycles gel-embedding-expansion treated mouse cerebellum (j). The MSI images were obtained with mass error tolerance of 10 ppm.

**Extended Data Figure 4.**
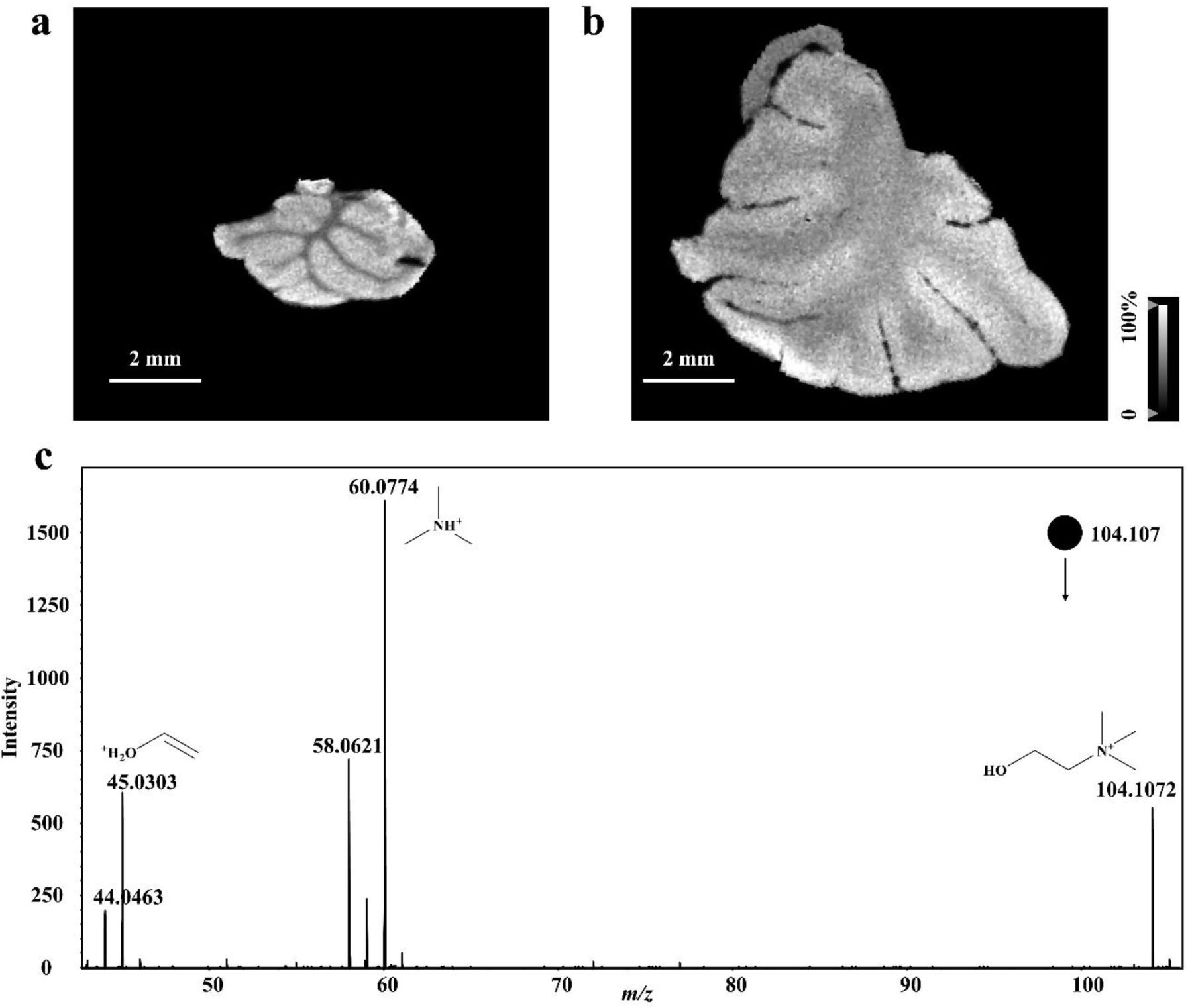
Gray MS images of choline (*m/z* 104.1072) from the unexpanded control and double-embedded cerebellum under positive mode. a. MS images of choline from the unexpanded control (MSI); b. MS images of choline from double-embedded cerebellum (TEMI); c. *In situ* MALDI-MS/MS of target ion of *m/z* 104.1072 from the expanded mouse cerebellum, which confirms the identification of Choline.

**Extended Data Figure 5.**
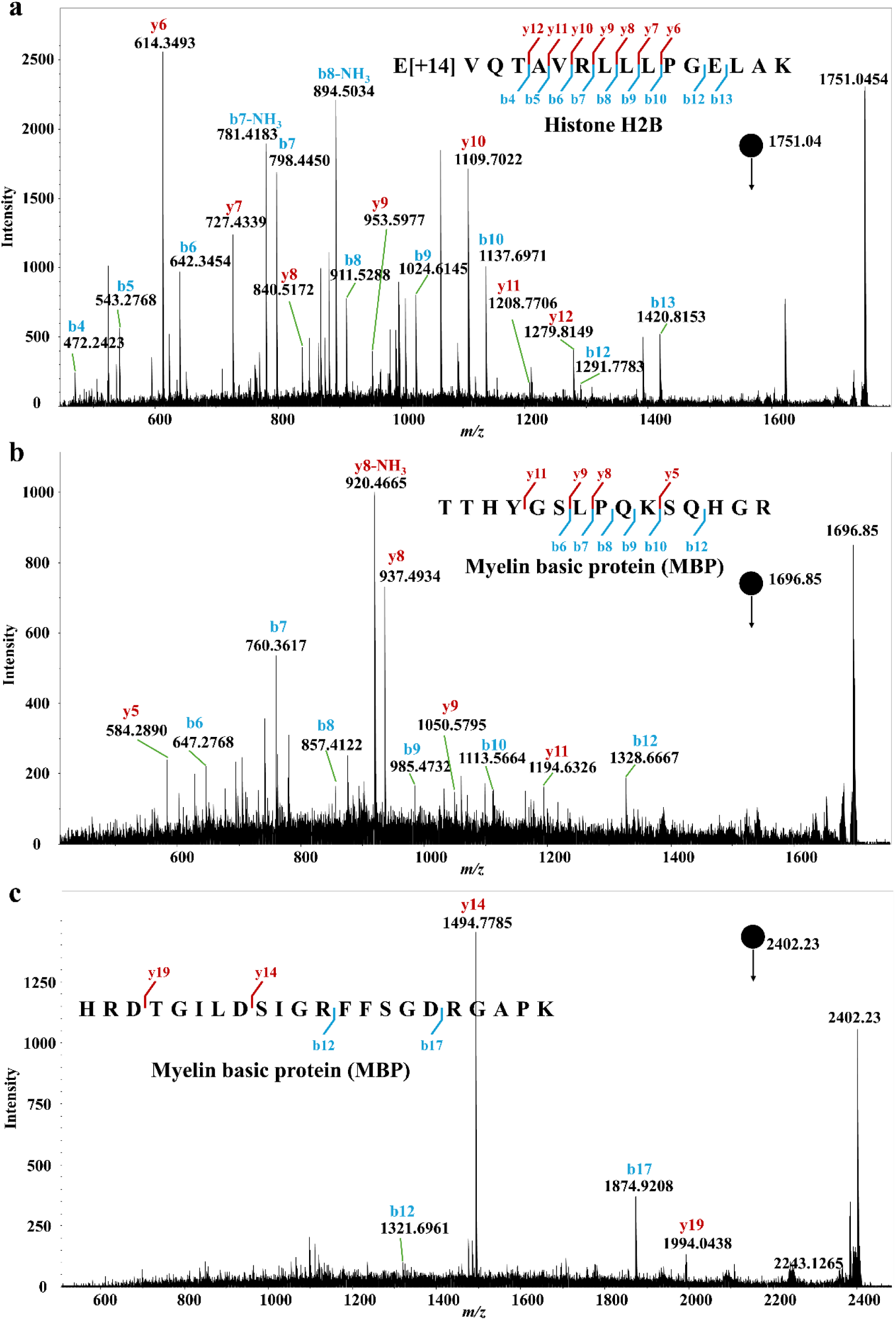
Representative tandem MS spectra of on-tissue tryptic peptides validation via *in situ* MALDI-MS/MS on the expanded mouse cerebellum. Annotated MALDI-MS/MS spectrum of precursor *m/z* 1751.04 that was acquired directly from the expanded tissue and identified as peptide E[+14]VQTAVRLLLPGELAK from Histone H2B, the E[+14] modification is caused by the Formaldehyde fixing; b. Annotated MALDI-MS/MS spectrum of precursor *m/z* 1696.85 and identified as peptide TTHYGSLPQKSQHGR from MBP; c. Annotated MALDI-MS/MS spectrum of precursor *m/z* 2402.23 and identified as peptide HRDTGILDSIGRFFSGDRGAPK from MBP.

## Materials and Methods

### Chemicals and materials

Reagents including Optima^TM^ LC/MS grade methanol (MeOH), ethanol (EtOH), acetonitrile (ACN), 2-propanol (IPA), water, and formic acid were purchased from Fisher Scientific (Pittsburgh, PA). 6-((acryloyl)amino) hexanoic acid succinimidyl ester (AcX), sodium acrylate, acrylamide, N,N’-methylenebis acrylamide (Bis), N,N,N’,N’-Tetramethyl ethylenediamine (TEMED), ammonium persulfate (APS), 4-hydroxy TEMPO, 1,5-diaminonaphthalene (DAN), 2,5-Dihydroxybenzoic acid (DHB), N-(1-Naphthyl)ethylenediamine dihydrochloride (NEDC), DAPI (4’,6-Diamidino-2-Phenylindole, Ethylenediaminetetraacetic acid (EDTA), Triphenylphosphine (TPP), Butylated hydroxytoluene (BHT), *tert*-Butyl methyl ether (MTBE) (puriss. p.a., ≥99.5% (GC)), and LiChropur^TM^ Ammonium formate (Crystals, ≥99.0%, Eluent additive for LC-MS) were purchased from Sigma-Aldrich (St. Louis, MO). SPLASH® Lipidomix® Mass Spec Standard was purchased from Avanti Polar Lipids. Metabolomics QReSS^TM^ Standard Mix 1 was purchased from Cambridge Isotope Laboratories, Inc. Benchmark Scientific D1032-30 Prefilled 2.0 mL Tubes, Triple-Pure High Impact Zirconium Beads, 3.0 mm was purchased from Benchmark Scientific. Indium tin oxide (ITO)-coated glass microscope slides (25 mm × 75 mm × 1.1 mm) were purchased from Delta Technologies (Loveland, CO).

Animal experiments received approval from the Animal Care and Use Committee of the University of Wisconsin-Madison and Janelia Research Campus, Howard Hughes Medical Institute. Mouse brain samples were harvested from female Wild Type mice (aged 6 to 8 months, Wild Type mice C57BL/6). Mice were housed in facilities with a standard light-dark cycle, humidity of 50% at 24 °C. The mice were housed and treated under a protocol approved by the Institutional Animal Care and Use Committee (IACUC) at the University of Wisconsin–Madison or Howard Hughes Medical Institute (Janelia Research Campus).

For collection of fixed mouse brain tissues, animals were transcardially perfused with 4% formaldehyde in 1× PBS, followed by post-fixation of the brain in the same solution overnight at 4 °C. Fixed tissue was washed in 1× PBS and sectioned by vibratome. For collection of fresh mouse brain tissues, euthanasia by decapitation was performed, and the whole brain tissue was surgically collected and snap frozen in liquid nitrogen immediately. The mouse brain tissue samples were stored at −80 °C until use.

Mouse tumor samples were harvested from a murine melanoma model. For tumor experiments, female Wild Type C57BL/6 mice at age 10 to 14 weeks were purchased from Taconic Biosciences, Inc. (Rensselaer, NY). Murine melanoma B78-D14 (B78) cell line, supplied by ATCC, was grown at 37 °C with 5% CO_2_ in RPMI-1640 medium (Gibco) containing 10% fetal bovine serum (FBS) (Gibco) and 1% penicillin–streptomycin solution (Gibco). B78 tumors were engrafted by subcutaneous flank injection of 4 × 10^6^ murine melanoma B78-D14 (B78) cells. Tumor size was determined using calipers and volume approximated as (width^2^ × length)/2 and the tumor samples were harvested at size ∼600 mm^3^, which was done approximately 5 weeks following the engraftment treatment.

Fresh mouse brain and tumor samples were fixed in 4% formaldehyde 1× PBS for 4 hours at room temperature, followed by washing with 1× PBS for three times. Then the tissue blocks were embedded in 6% (wt/wt) agarose and sectioned into tissue slices at 500 *µ*m thickness using a vibratome (Leica VT1000 S).

### Sample preparation for TEMI

The schematic workflow of sample preparation for TEMI is presented in **Figure 1**. Tissue expansion was initially based on prior implementations of expansion microscopy^23^, however, adjustments have been made to tailor the expanded samples specifically for MALDI-MS imaging purposes. Mice were perfused with formaldehyde as described above, organs dissected, and the adjacent tissue slices were cut into 500 *µ*m and 300 *µ*m for unexpanded control and for TEMI sample preparation respectively.

For anchoring proteins prior to gelation, the 300 *µ*m tissue slice was treated in 0.2 mg/mL AcX in 1× PBS solution for 1 h at room temperature, followed by washing with 1× PBS. Stock solutions of monomer solution components were prepared and stored frozen as in our previous study^15^: 40% acrylamide, 4 M sodium acrylate, 1% (w/v) bisacrylamide, 0.5% (w/v) 4-hydroxy TEMPO (4HT), 10% (v/v) TEMED, and 10% (w/v) APS in deionized water. TREx1000 gelation solution was prepared according to the recipe from our previous study^15^ with final concentrations of 1 M sodium acrylate, 14% acrylamide, 0.1% bisacrylamide, 1× PBS, 0.01 % 4HT, 0.2% TEMED, and 0.2% APS. All components but 4HT, TEMED, and APS were mixed to produce monomer solution, aliquoted, and stored frozen. 4HT, TEMED, and APS (gelation inhibitor, accelerator, and initiator) stocks were thawed and added to monomer solution on ice to produce complete gelation solution within minutes of use. Tissue slices were incubated in complete gelation solution on ice for 30 minutes with shaking before being mounted in gelation chambers and gelled in a humidified 37 °C incubator for 2 h as previously described^15^.

After gelation, the gelation chamber was carefully disassembled, and the gelled tissue was trimmed out with a scalpel. Then the gelled tissue was collected and incubated in 1× PBS for 30 min at 4 °C and re-embedded in the same gelation solution for each additional round of gelation. After the desired rounds of embedding-expansion, both unexpanded and expanded tissues were optionally stained with 0.5 μg/mL DAPI and/or 1 μg/mL Alexa488 NHS ester dye in 1× PBS for overnight or 1 h followed by washing with 1× PBS, 3× 30 min, for fluorescence imaging of DNA and/or proteins, respectively. For different lumiprobe staining is advised by the corresponding application with MSI, for example, if one is interested in detecting DNA, DAPI staining should be skipped; if one is interested in detecting peptides or proteins, Alexa488 NHS ester dye staining should be skipped as it occupies the primary amines of lysine and N-termini of proteins.

Both unexpanded and expanded tissues were incubated in 30% (w/w) sucrose cryoprotectant in 1× PBS for 4 h (the sucrose solution was replaced with new solution every hour), to protect against the formation of large ice crystals during cryosectioning. Then, the gel was embedded in 10% (w/w) gelatin after trimming away extra gel outside the tissue, gently pressed flat in case of mild curvature and snap-frozen on dry ice prior to cryosectioning at 30 *µ*m in a cryostat (Thermo Scientific) at −30 °C. Tissue sections were mounted on poly-L lysine (Electron Microscopy Sciences) coated ITO slides and dried in a desiccator for 20 min, and then stored in −80 °C until MSI processing.

The tissue slides were thawed and dried again in a desiccator for 30 min prior to the application of MALDI matrix. For dual-polarity MS imaging of lipid species, DAN matrix (10 mg/mL) in acetonitrile/water (v/v, 80:20) solution was sprayed onto the tissue samples at a flow rate of 75 *μ*L/min with tracking space of 3 mm for 6 passes using a M5-Sprayer (HTX Technologies, Carrboro, NC, USA). The moving velocity of the nozzle was set to 1000 mm/min, the nozzle temperature was set to 75 °C, and the nozzle nitrogen gas pressure was 10 psi. A drying time of 30 sec was applied between each pass. For small metabolites, N-glycan, and tryptic peptide analyses in positive ionization mode, DHB (40 mg/mL with 0.1% formic acid in acetonitrile/water (v/v, 80:20) solution) was used under the same parameters setting of the M5-Sprayer; while for small metabolite analysis in negative mode, NEDC matrix (10 mg/mL with 0.2% ammonium hydroxide in acetonitrile/water (v/v, 80:20) solution) was used with the same M5-Sprayer settings.

For N-glycan and tryptic peptide MS imaging on the same tissue slides following prior lipid or metabolite analysis, the remaining MALDI matrix on the tissue was removed by washing three times in 100% ethanol, 95% ethanol and 70% ethanol for 5 minutes per wash. Then the tissue slides were dried in a fume hood at room temperature, followed by immersion in an antigen retrieval buffer containing 100 mM acidic Tris and 1 M hydroxylamine hydrochloride for 2 hours at 60 °C. A 10% (v/v) TFA solution was used to adjust pH of the antigen retrieval buffer to 3. After the decrosslinking-inducing antigen retrieval treatment, the slides were washed three times with 95% and 70% ethanol solutions for 2 minutes, respectively.

For *in situ* releasing of N-glycans, a 20 μL of PNGase F (20 U, Promega) was dissolved in 380 *μ*L of 20 mM ammonium bicarbonate (ABC) buffer, which was sprayed onto tissue sections via the TM-Sprayer with the following setup: flow rate of 20 *μ*L/min, nitrogen gas pressure at 8 psi, nozzle temperature at 35°C, nozzle velocity of 800 mm/min, and tracking space of 3 mm. Following incubation in a humid chamber at 37 °C for 8 h, sections were sprayed with DHB (40 mg/mL with 0.1% formic acid in acetonitrile/water (v/v, 80:20)) using the TM-Sprayer with the aforementioned DHB matrix spraying settings.

For label-free protein MS imaging, *in situ* tryptic digestion was performed. Briefly, tissue sections were washed with 70% ethanol to remove the remaining DHB matrix and released N-glycans, placed in fume hood until dry, then sprayed with freshly prepared 50 ng/μL trypsin (Promega) in 20 mM ABC buffer using the same TM-Sprayer parameters described above for PNGase F deposition. After incubation in a humid chamber at 37°C for 8 h, sections were sprayed with DHB (40 mg/mL with 0.1% formic acid in acetonitrile/water (v/v, 80:20)) using TM-Sprayer with the same TM-Sprayer parameters described above for matrix application.

For antibody-based targeted protein MSI, photocleavable mass-tag (PC-MT) MALDI-immunohistochemistry (IHC) was performed. PC-MT tagged antibody (AmberGen) staining was based on AmberGen’s protocol^24^. The working antibody probes were diluted in tissue blocking buffer (2% (v/v) Normal Rabbit Serum, 2% (v/v) Normal Mouse Serum, and 5% (w/v) BSA prepared in TBS (50 mM Tris-HCl, pH 7.5, 200 mM NaCl) with 0.05% (w/v) Octyl β-D-Glucopyranoside) to a final concentration of 2.5 μg/mL. The PC-MT probes on the stained tissue sections were photocleaved by illumination with 365 nm UV light with a Phrozen UV curing lamp for 10 min (3 mW/cm^2^) to achieve maximum photocleavage prior to applying α-Cyano-4-hydroxycinnamic acid (CHCA) matrix (10 mg/mL with 0.1% formic acid in acetonitrile/water (v/v, 80:20) solution) on the samples. TM-Sprayer settings are implemented as below: flow rate of 100 *μ*L/min for 6 passes, drying time of 30 s between each pass, nozzle nitrogen gas pressure at 10 psi, nozzle temperature at 75 °C, nozzle velocity of 1000 mm/min, and tracking space of 3 mm.

### Extraction of mouse brain lipidome for LC-MS analysis

Cerebellar slices were cut with a thickness of 300 *µ*m. Three groups, control (without expansion), Ex (expanded without applying proteinase K), and ExP (expanded with applying proteinase K) were used for lipidomics analysis. The wet mass of slices was around 5 mg each for unexpanded controls, or 70 mg after expansion. All the slices were washed in ddH_2_O 2× 1.5 h in a bead-beater tube. After removing all the water, 200 *µ*L of LC-MS water with 5 *µ*L antioxidant stock solution (0.2 mg/mL of TPP, 0.2 mg/mL of BHT and 1 mg/mL of EDTA) was added. Then the sample was homogenized with 3.0 mm zirconium beads at the highest speed for 3 min (6 cycles of 30 sec inter-rest). Then all the homogenized solutions were transferred into a 4-mL wide-mouth glass vial and added 5 µL of SPLASH® LIPIDOMIX® Mass Spec Standard. Then the lipids were extracted by adding methanol (LC-MS grade) and vortex for 1 min, followed by adding MTBE and shaking for 1 h at 25 °C, and then the phase separation was induced by adding LC-MS-grade water with a final mixing ratio of MTBE/methanol/water (10:3:2.5, v/v/v)^25^. Note that since the expanded sample was swollen with water by the hydrogel, there was no extra water needed for these specimens. The upper (organic) phase was collected and dried under nitrogen gas stream. The lipid extracts were stored at −80 °C before analysis.

### Extraction of polar metabolome from mouse cerebellum

The polar metabolites were extracted from the mouse cerebellum by dry-ice cold 80% methanol. The sample, mixed with 2 *µ*L of Metabolomics QReSS^TM^ Standard Mix 1 followed by the manufacture’s instruction, was homogenized with 3.0 mm zirconium beads at the highest speed for 6 cycles of 30 sec with 5 min cool down in between the rounds.

### Extraction of tryptic peptides for LC-MS/MS analysis

The tryptic peptides on the tissue sections were extracted with 10 *µ*L of 50% acidic ACN (0.1% FA) and 80% acidic ACN (0.1% FA) for five times, respectively. Briefly, an aliquot of 10 *µ*L solvent were deposited onto the tissue section using a pipette, after 3-sec extraction, the solvent was retracted back into the pipette and transferred into a 0.6 mL vial. All the peptide extraction solutions from three tissue section replicates were combined and dried in a SpeedVac. The samples were reconstituted in 100 *µ*L 0.1% FA water and subjected to desalting using OMIX C18 Pipette Tips (Agilent). The process began by wetting the tip with three aspirating cycles of acetonitrile (ACN). Next, the tip was equilibrated by aspirating a 0.1% formic acid water three times. Following this, a 90 *µ*L sample volume was aspirated into the pipette tip and cycled at least ten times for thorough enrichment. The tip was then washed three times with the 0.1% formic acid water solution. Two clean microcentrifuge tubes were prepared, each containing 100 *µ*L of 50% and 80% ACN water mixture (v/v) with 0.1% formic acid. Use the solutions to aspirate and dispense eluant through the C18 pipette tip, cycling at least ten times to ensure effective elution of the desalted peptides. Finally, the two eluents were combined and dried *in vacuo*.

### Data acquisition and analysis

All the MALDI-MSI experiments were carried out using a timsTOF fleX mass spectrometer (Bruker Daltonics) with timsContol and flexImaging for data acquisition, which was coupled with a SmartBeam 3D 10 kHz frequency tripled Nd:YAG laser (355 nm). The laser settings used were 10-50 *μ*m diameter at smart mode, with 100 shots per pixel and a raster step size that corresponded to the laser spot size during the MS imaging. The laser power was set to 40-80% with laser shots of 100-250. For lipid analysis, the MS imaging data were collected under positive ionization mode over a mass range of 400-1000 Da; for the subsequent negative imaging run on the same tissue sample, 100 shots with laser energy of 45% were conducted. It is important to note that, in dual ionization of the same tissue section, it is crucial to carefully control the laser energy at an appropriate level and refrain from using excessive laser energy during the first MSI run, as excessive laser energy could damage the tissue and ultimately compromise the performance in the subsequent MSI run. A varied mass range of 100–3000 Da was used for imaging small metabolites, tryptic peptides, and N-glycans. The mass resolving power of the instrument is typically 40000 (fwhm) in the lipid mass range, resulting in high mass accuracy in MS imaging experiments. Other instrumental parameters include collision RF of 800–4000 Vpp, ion transfer time of 20–80 *μ*s, prepulse storage time of 5–10 *μ*s, and multipole RF of 350 Vpp. In MALDI-MS/MS experiments, target precursor lipid ions were isolated at a mass window of 1.5 Da for HCD fragmentation with collision voltage of 30-60 eV. MS data were analyzed with Compass Data Analysis (Bruker) and the MSI images were visualized using SCiLS Lab Pro (Bruker) with data normalized to total ion count (TIC) and no denoising processed. Unsupervised statistical analysis tool of Bisecting k-means was used for an arbitrarily segmentation of all single spectrum from the sample based on the detected peaks of lipids; each segment is annotated with a different color. Lipids, small metabolites, and tryptic peptides detected from the tissue samples were identified based on the combination of *in situ* MALDI-MS/MS of target analytes, as well as referring to high performance liquid chromatography (HPLC)-nanoESI-MS/MS results of metabolite, lipid, and tryptic peptide extracts from the tissue samples. The compositions of N-glycans observed were tentatively assigned through searches in the GlyGen (http://www.glygen.org) and GlycoWorkbench (https://code.google.com/archive/p/glycoworkbench/) with a mass error tolerance of 10 ppm.

For conventional lipidomic analysis, the lipid extracts were re-dissolved in 55 *µ*L of IPA:MeOH (1:1) and 12 *µ*L was injected to run LC-MS. The experiments were performed on an Orbitrap Fusion Lumos Tribrid Mass Spectrometer equipped with Ion Max API source housing with HESI-II probe and Vanquish UHPLC System (ThermoFisher Scientific). The analytes were separated on a Accucore^TM^ Vanquish^TM^ C18+ UHPLC Column (2.1 x 150 mm, 1.5 *µ*m) column operated at 55 °C and a flow rate of 150 *µ*L/min using a 52 min gradient. The gradient was shown in the **Supplementary Table 7**. The mobile phase A was 60:40 (v:v) acetonitrile/water with 10 mM ammonium formate and 0.1% formic acid and mobile phase B was 90:10 (v:v) IPA/acetonitrile with 10 mM ammonium formate and 0.1% formic acid. The MS spectra were acquired with Xcalibur software (ThermoFisher Scientific). The comprehensive data dependent HCD MS2 experiment with conditional CID MS2 and MS3 data acquisition was conducted. Alternating positive and negative ion data-dependent HCD MS2 experiments were performed using two 1.0 s cycles (0–31.2 min retention time range) with stepped collision energy (24%, 27%, 30%). Additional CID MS2 scans were triggered on the same precursor ion for PC lipids with a diagnostic fragment ion (*m/z* 184.0733) detected from the positive HCD MS2 data. For retention time ranging 31.2–52 min, a positive ion, data-dependent MS2 experiment was performed using a 2 s cycle time. Additional CID MS3 scans were subsequently triggered on the three largest HCD product ions that lost neutral fatty acid plus ammonia.

The raw files were analyzed by LipidSearch5 (Thermo Fisher Scientific). The lipids were identified by matching product ion spectra to the LipidSearch library (**Supplementary Data 1**). Both precursor and product ion mass error tolerance were set at 5 ppm and aligned based on retention time tolerance of 0.05 min. After manually removing the duplicates and checking the peak and product ions, as well as selecting the main ion for each lipid, the final dataset with 659 lipids was used for further analysis. The positive ions were normalized to the internal reference of PE (18:1_15:0) (d7) in each sample, and the negative ions were normalized to the internal reference of PC (15:0_18:1) (d7) in each sample. After normalization, all the data were grouped and filtered based on at least 3 valid values in at least one group. The data was log_2_-transformed, and the missing values were imputed based on normal distribution. The ANOVA test was performed for significance analysis, followed by post hoc Tukey’s HSD test (**Supplementary Table 8**). The Benjamini-Hochberg adjusted p value that less than or equal to 0.05 was considered as significance using Perseus^26, 27^.

For LC-MS/MS analysis of polar metabolites, an Orbitrap Fusion Lumos Tribrid Mass Spectrometer equipped with Ion Max API source housing with HESI-II probe and Vanquish UHPLC System (ThermoFisher Scientific) was used. Four microliters of the resuspended sample in methanol were injected onto a Waters Acquity UPLC BEH Amide column (150 mm× 2.1 mm, 1.7 *µ*m) operated at 45 °C and a flow rate of 200 *µ*L/min using a 68 min gradient. The gradient was shown in the **Supplementary Table 9**. The mobile phase A was water with 10 mM ammonium formate and 0.125% formic acid, the mobile phase B was made from acetonitrile/water (95/5, v/v) with 10 mM ammonium formate and 0.125% formic acid. The ion source conditions were set as follows: spray voltage: 3.5 kV for positive mode, 2.5 kV for negative mode, sheath gas: 50 arbitrary units, aux gas: 10 arbitrary units, sweep gas: 1 arbitrary unit, ion transfer tube temp, 325 °C; vaporizer temp, 350 °C. The following acquisition parameters were used for MS1 analysis: resolution: 120K, AGC target: standard, Maximum IT: auto, scan range: 60–700 *m/z*, Data type: profile. Data-dependent MS/MS parameters: resolution: 30K, AGC target: standard, Maximum IT: auto, isolation window: 0.7 *m/z*, Activation type: HCD, Collision energy mode: stepped: 20, 30, 40, Data type: centroid. The raw files are analyzed using Compound Discoverer (ThermoFisher Scientific) for untargetted metabolomics.

For LC-MS/MS analysis of extracted tryptic peptides, an Orbitrap Ascend Tribrid Mass Spectrometer equipped with Vanquish Neo UHPLC System (ThermoFisher Scientific) was used. Five microliters of the sample were injected onto an Aurora Ultimate XT 25×75 C18 UHPLC column (25 cm× 75 *µ*m, 1.7 *µ*m) operated at 35 °C and a flow rate of 200 nL/min using a 130 min gradient. The gradient was shown in the **Supplementary Table 10**. The mobile phase A was deionized water with 0.1% formic acid, the mobile phase B 80% acetonitrile with 0.1% formic acid. The ion source conditions were set as follows: spray voltage: 1.9 kV at positive mode, ion transfer tube temp, 275°C. The following acquisition parameters were used for MS1 analysis: resolution: 120K, normalized AGC target: 250%, Maximum IT: 50 ms, scan range: 300–2000 *m/z*, Data type: profile. Data-dependent MS/MS parameters: resolution: 15K, AGC target: standard, Maximum IT: auto, isolation window: 1.1 *m/z*, Activation type: HCD, Collision energy mode: normalized: 25%, Data type: centroid.

The raw files were analyzed by FragPipe based MSFragger. The tryptic peptides were searched against a database containing reviewed proteome for *Mus musculus* database from Uniprot with false discovery rate for precursors set at 1%. The precursor mass tolerance was set to 10 ppm, maximum of four missed cleavages was allowed. Dynamic modifications including deamidation of asparagine and glutamine, oxidation on methionine and cysteine, and +12 Da, +14.017 Da, +30.011 Da on any amino acids was allowed.

### Expansion deformation mapping

To measure non-uniformity of expansion, we stained the tissue with 0.5 *μ*g/mL Alexa488 NHS ester dye after the anchoring step and imaged at each stage of gel embedding. We analyzed the images using BigWarp (Fiji) and MATLAB as described previously^28, 29^. Briefly, the expanded tissue image was registered to the pre-expansion tissue image using manually picked landmarks. The deformation field produced from these landmarks was used to calculate the expansion factor and measure the degree of deformation due to expansion non-uniformity (**Supplementary Note 2**).

## Data Availability

Source data is provided with this paper. The MS imaging data included in this study are available from the corresponding author upon request. All the liquid chromatography-mass spectrometry raw files have been deposited into MassIVE (dataset identifier MSV000096466, ftp://massive.ucsd.edu/v07/MSV000096466/). All the Source Data for Figures 1-6 and Extended Data Figures 3 and 4 of MALDI-MSI raw files have been deposited into MassIVE (dataset identifier MSV000097036, ftp://massive.ucsd.edu/v09/MSV000097036/).

## Code Availability

The code of data for the quantification of tissue expansion non-uniformity is available in Supporting Information materials as **Supplementary Note 3**.

## Supplementary Information

Supplementary Information

Supplementary Figures 1–18, Supplementary Tables 1–10, and Supplementary Notes 1–3.

Supplementary Data 1

## References

1. Rao, A., Barkley, D., Franca, G.S. & Yanai, I. Exploring tissue architecture using spatial transcriptomics. Nature 596, 211–220 (2021).

2. Tian, L.Y., Chen, F. & Macosko, E.Z. The expanding vistas of spatial transcriptomics. Nat. Biotechnol. 41, 773–782 (2023).

3. Capolupo, L. et al. Sphingolipids control dermal fibroblast heterogeneity. Science 376, eabh1623 (2022).

4. Buchberger, A.R., DeLaney, K., Johnson, J. & Li, L.J. Mass Spectrometry Imaging: A Review of Emerging Advancements and Future Insights. Anal. Chem. 90, 240–265 (2018).

5. Vicari, M. et al. Spatial multimodal analysis of transcriptomes and metabolomes in tissues. Nat. Biotechnol. (2023).

6. Zhang, H., Delafield, D.G. & Li, L.J. Mass spectrometry imaging: the rise of spatially resolved single-cell omics. Nat. Methods 20, 327–330 (2023).

7. Schulz, S., Becker, M., Groseclose, M.R., Schadt, S. & Hopf, C. Advanced MALDI mass spectrometry imaging in pharmaceutical research and drug development. Curr. Opin. Biotechnol. 55, 51–59 (2019).

8. Yuan, Z.Y. et al. SEAM is a spatial single nuclear metabolomics method for dissecting tissue microenvironment. Nat. Methods 18, 1223–1232 (2021).

9. Niehaus, M., Soltwisch, J., Belov, M.E. & Dreisewerd, K. Transmission-mode MALDI-2 mass spectrometry imaging of cells and tissues at subcellular resolution. Nat. Methods 16, 925–931 (2019).

10. Bien, T., Koerfer, K., Schwenzfeier, J., Dreisewerd, K. & Soltwisch, J. Mass spectrometry imaging to explore molecular heterogeneity in cell culture. Proc. Natl. Acad. Sci. U. S. A. 119, e2114365119 (2022).

11. Passarelli, M.K. et al. The 3D OrbiSIMS-label-free metabolic imaging with subcellular lateral resolution and high mass-resolving power. Nat. Methods 14, 1175–1183 (2017).

12. Kompauer, M., Heiles, S. & Spengler, B. Atmospheric pressure MALDI mass spectrometry imaging of tissues and cells at 1.4-mu m lateral resolution. Nat. Methods 14, 90–96 (2017).

13. Chen, F., Tillberg, P.W. & Boyden, E.S. Expansion microscopy. Science 347, 543–548 (2015).

14. Truckenbrodt, S., Sommer, C., Rizzoli, S.O. & Danzl, J.G. A practical guide to optimization in X10 expansion microscopy. Nature Protocols 14, 832–863 (2019).

15. Damstra, H.G.J. et al. Visualizing cellular and tissue ultrastructure using Ten-fold Robust Expansion Microscopy (TREx). Elife 11, e73775 (2022).

16. Chang, J.B., et al. Iterative expansion microscopy. Nat. Methods 14, 593–599 (2017).

17. Chen, L.C., Lee, C.P. & Hsu, C.C. Towards developing a matrix-assisted laser desorption/ionization mass spectrometry imaging (MALDI MSI) compatible tissue expansion protocol. Anal. Chim. Acta 1297 (2024).

18. Hung, Y.L.W. et al. Expansion Strategy-Driven Micron-Level Resolution Mass Spectrometry Imaging of Lipids in Mouse Brain Tissue. CCS Chemistry, 1–9 (2024).

19. Chan, Y.H. et al. Gel-assisted mass spectrometry imaging enables sub-micrometer spatial lipidomics. Nat. Commun. 5036 (2024).

20. Samuel, J.M., Yan, T., Liang, Z. & Prentice, B.M. Examination of Lipid Distributions in Hydrogel-Expanded Mouse Brain Tissue Using Imaging Mass Spectrometry. bioRxiv (2024).

21. Xie, C., et al. Ten-Fold Expansion MALDI Mass Spectrometry Imaging of Tissues and Cells at 500 nm Resolution. bioRxiv (2024).

22. Monroe, E.B. et al. Massively parallel sample preparation for the MALDI MS analyses of tissues. Anal. Chem. 78, 6826–6832 (2006).

## Methods-only references

23. Tillberg, P.W. et al. Protein-retention expansion microscopy of cells and tissues labeled using standard fluorescent proteins and antibodies. Nat. Biotechnol. 34, 987–992 (2016).

24. Yagnik, G., Liu, Z.Y., Rothschild, K.J. & Lim, M.J. Highly Multiplexed Immunohistochemical MALDI-MS Imaging of Biomarkers in Tissues. J. Am. Soc. Mass. Spectrom. 32, 977–988 (2021).

25. Matyash, V., Liebisch, G., Kurzchalia, T.V., Shevchenko, A. & Schwudke, D. Lipid extraction by methyl-tert-butyl ether for high-throughput lipidomics. J. Lipid Res. 49, 1137–1146 (2008).

26. Del Prete, E. et al. ADViSELipidomics: a workflow for analyzing lipidomics data. Bioinformatics 38, 5460–5462 (2022).

27. Tyanova, S. et al. The Perseus computational platform for comprehensive analysis of (prote)omics data. Nat. Methods 13, 731–740 (2016).

28. Jurriens, D., van Batenburg, V., Katrukha, E.A. & Kapitein, L.C. in Expansion Microscopy for Cell Biology, Vol. 161. (eds. P. Guichard & V. Hamel) 105–124 (2021).

29. Damstra, H.G. et al. GelMap: intrinsic calibration and deformation mapping for expansion microscopy. Nat. Methods 20, 1573–1580 (2023).

